# High Volume Rate 3-D Ultrasound Imaging Using Fast-Tilting and Redirecting Reflectors

**DOI:** 10.1101/2023.03.07.531439

**Authors:** Zhijie Dong, Shuangliang Li, Xiaoyu Duan, Matthew R. Lowerison, Chengwu Huang, Qi You, Shigao Chen, Jun Zou, Pengfei Song

**Affiliations:** Beckman Institute for Advanced Science and Technology, University of Illinois Urbana-Champaign, Urbana, IL; Department of Electrical and Computer Engineering, University of Illinois Urbana-Champaign, Urbana, IL; Department of Electrical and Computer Engineering, Texas A&M University, College Station, TX; Department of Radiology, Mayo Clinic College of Medicine and Science, Rochester, MN; Department of Bioengineering, University of Illinois Urbana-Champaign, Urbana, IL

**Author notes:** Corresponding Author: Dr. Pengfei Song, Department of Electrical and Computer Engineering, Beckman Institute for Advanced Science and Technology, University of Illinois Urbana-Champaign, 405 N. Mathews Ave., Urbana, IL 61801.

## Abstract

3-D ultrasound imaging has many advantages over 2-D imaging such as more comprehensive tissue evaluation and less operator dependence. Although many 3-D ultrasound imaging techniques have been developed in the last several decades, a low-cost and accessible solution with high imaging volume rate and imaging quality remains elusive. Recently we proposed a new, high volume rate 3-D ultrasound imaging technique: Fast Acoustic Steering via Tilting Electromechanical Reflectors (FASTER), which uses a water-immersible and fast-tilting acoustic reflector to steer ultrafast plane waves in the elevational direction to achieve high volume rate 3-D ultrasound imaging with conventional 1-D array transducers. However, the initial implementation of FASTER imaging only involves a single fast-tilting acoustic reflector, which is inconvenient to use because the probe cannot be held in the regular upright position. Also, conventional FASTER imaging can only be performed inside a water tank because of the necessity of using water for acoustic conduction. To address these limitations of conventional FASTER, here we developed a novel ultrasound probe clip-on device that encloses a fast-tilting reflector, a redirecting reflector, and an acoustic wave conduction medium. The new FASTER 3-D imaging device can be easily attached to or removed from clinical ultrasound transducers, allowing rapid transformation from 2-D to 3-D ultrasound imaging. *In vitro* B-mode imaging studies demonstrated that the proposed method provided comparable imaging quality (e.g., spatial resolution and contrast-to-noise ratio) to conventional, mechanical-translation-based 3-D imaging while providing a much faster 3-D volume rate (e.g., 300 Hz vs ∼10 Hz). In addition to B-mode imaging, we also demonstrated 3-D power Doppler imaging and 3-D super-resolution ultrasound localization microscopy with the newly developed FASTER device. An *in vivo* imaging study showed that the FASTER device could clearly visualize the 3-D anatomy of the basilic vein of a healthy volunteer, and customized beamforming was implemented to accommodate the speed of sound difference between the acoustic medium and the imaging object (e.g., soft tissue). These results suggest that the newly developed redirecting reflector and the clip-on device could overcome key hurdles for future clinical translation of the FASTER 3-D imaging technology.

## Introduction

Ultrasound is a widely available and accessible medical imaging modality thanks to its safety, real-time imaging speed, portability, and low cost. In the last two decades, driven by the rapid growth of ultrafast imaging technologies [1-3], many advanced ultrasound imaging techniques such as shear wave elastography (SWE) [4], functional ultrasound (fUS) [5], and super-resolution ultrasound localization microscopy (ULM) [6] have been rapidly emerging. These imaging techniques provided new biomarkers that created numerous unprecedented possibilities in many preclinical and clinical applications [7].

Despite the rapid development of ultrasound imaging, the majority of ultrasound scans being performed are still 2-D. This forms one of the key challenges of ultrasound imaging because the 2-D scanning is operator dependent and may not provide comprehensive evaluation of the targeted tissue. Furthermore, one of the major functions of ultrasound imaging is to detect motions (e.g., blood flow, shear wave, and microbubble flow), which propagate in all three dimensions, and cannot be fully captured with 2-D scans. This results in inaccurate quantifications of biomarkers such as blood flow velocity and shear wave speed if the blood flow and shear wave motion have out-of-plane components [8, 9]. Moreover, 3-D ultrasound scanning is also beneficial for many intraoperative and interventional procedures that use ultrasound images for guidance [10-13].

Although 3-D ultrasound imaging systems are present in the clinic for echocardiography [14-16], surgical guidance [17], obstetrics [15], and vascular imaging [18], their performance and functionality are limited by high equipment costs (e.g., 2-D matrix arrays), being cumbersome to use (e.g., wobblers), and low scanning volume rates. For example, the most popular clinical 3-D ultrasound solution is based on “wobblers” that mechanically sweep a 1-D probe to collect 2-D imaging slices that are stacked into 3-D volumes [19]. However, this inevitably results in bulky devices and is limited to very slow scanning speed (e.g., several volumes per second). Another 3-D imaging solution is based on 2-D matrix arrays, which support electronic scanning of large 3-D volumes at a much higher volume rate (e.g., tens of Hertz). However, 2-D matrix arrays are expensive and only available on high-end ultrasound systems for specialty applications in the clinic. Furthermore, due to a large number of transducer elements and channels, 2-D matrix arrays are also associated with high computational costs (e.g., beamforming). As such, multiplexing [20] or micro-beamforming [21] are typically necessary for practical implementations of 2-D matrix arrays. Nevertheless, the 3-D volume rate is compromised due to the need for multiple pulse-echo cycles to reconstruct a full 3-D volume. Although emerging techniques such as sparse arrays [22] and row-column addressing (RCA) arrays [23-26] present promising solutions for high volume-rate 3-D scanning, each of them has its own challenges and still requires further development.

Recently, we have investigated a new ultrasound scanning technique, called Fast Acoustic Steering via Tilting Electromechanical Reflectors (FASTER) [27], which uses a water-immersible and fast-tilting acoustic reflector to achieve low-cost and high volume-rate 3-D scanning by rapidly steering ultrafast plane waves in the elevational direction. FASTER 3-D imaging does not need 2-D matrix arrays and can be directly applied to conventional 1-D array transducers. Unlike the wobbler, FASTER sweeps the ultrasound beam instead of the ultrasound transducer, thereby enabling a much faster 3-D scanning speed (e.g., hundreds of Hertz). Another advantage of FASTER over wobblers is that because FASTER does not sweep the transducer, it does not use a motor and thus can have a much more compact and lightweight form factor. However, one limitation was that since the fast-tilting reflector alters the ultrasound wave propagation direction (i.e., from axial to elevational propagation), the ultrasound probe could not be held in the regular position. In this study, a second acoustic reflector (i.e., a redirecting reflector) was added to recover the axial acoustic wave propagation direction, which allows the transducer to be held in a regular upright position by the operator. In addition, a compact and lightweight clip-on housing device was designed and fabricated to enclose both reflectors and the acoustic wave coupling medium. As a result, the new FASTER clip-on device can be easily attached to or removed from conventional 1-D array transducers. Both *in vitro* and *in vivo* studies were conducted to evaluate the 3-D imaging capabilities of the new FASTER 3-D imaging device. The application of high volume rate 3-D scanning was also demonstrated with 3-D power doppler (PD) and 3-D ULM experiments.

The rest of this article is structured as follows. We first describe the design of the new FASTER 3-D clip-on device and related image reconstruction processes. We then present validation and calibration, followed by *in vitro* and *in vivo* imaging studies. We finalize the article with discussions and conclusions.

## Materials and Methods

### Principles of FASTER 3-D Imaging

The proposed FASTER technique takes advantage of the ultrafast frame rate of plane wave imaging [1] (e.g., thousands of Hertz to tens of thousands of Hertz) and rapidly distributes ultrasound beams at different elevational locations using a water-immersible and microfabricated fast-tilting reflector [27]. In the previous design, the imaging direction was orthogonal to the conventional axial imaging direction because the fast-tilting reflector bends the ultrasound beam by 90 degrees. To allow the transducer to be held in a regular position (e.g., upright), a redirecting reflector was added prior to the fast-tilting reflector to recover the axial incident beam direction (Fig. 1). The ultrasound beam is first reflected by the redirecting reflector by 90 degrees (from axial propagation to elevational), followed by another 90-degree reflection by the fast-tilting reflector (from elevational back to axial). The redirecting reflector and fast-tilting reflectors were made of the same silicon wafer, which provides high acoustic reflectivity due to the high acoustic impedance and smooth surface. The redirecting reflector was parallel to the stationary position of the fast-tilting reflector (i.e., tilted by 45 degrees to the incident beam direction), as shown in Fig. 1(c).

**Fig. 1.**
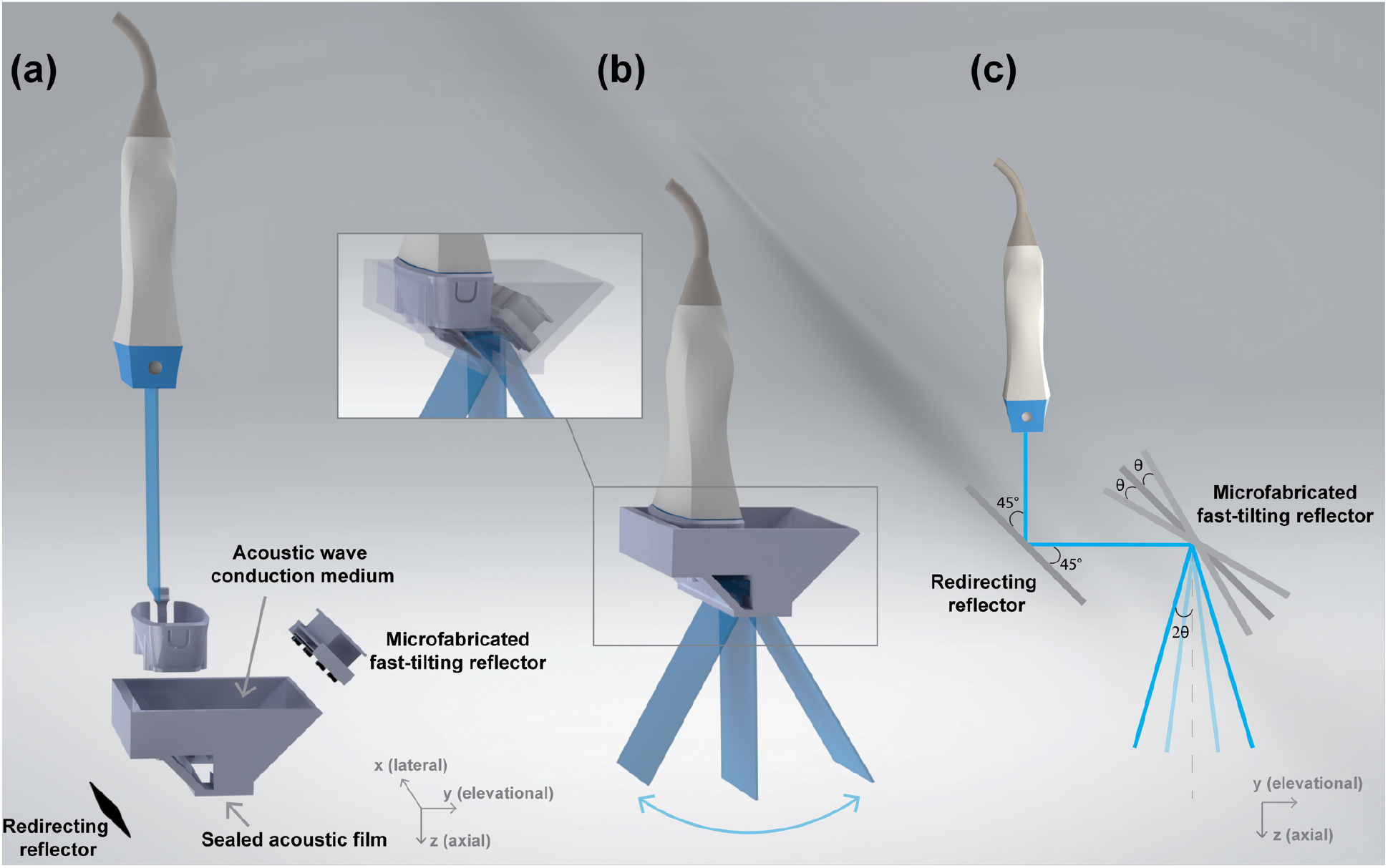
FASTER 3-D imaging device schematics. (a) Components of the new FASTER 3-D imaging device: conventional 1-D ultrasound probe and clip-on housing device, which encloses a redirecting reflector, a fast-tilting reflector (driven by two electromagnet coils), acoustic wave conduction medium, and an acoustic film (on the bottom) for sealing. (b) FASTER 3-D scanning schematics. The FASTER device rapidly distributes ultrasound beams at different elevational locations using the fast-tilting reflector. (c) Acoustic reflection path of the FASTER 3-D imaging device. The ultrasound beam is reflected by the redirecting reflector first by 90 degrees, followed by another reflection by the fast-tilting reflector.

To address the limitation of needing a water tank in the previous FASTER setup, a compact clip-on housing device was designed. The clip-on housing device encloses the reflectors and acoustic wave conduction medium (water at room temperature) with a sealing acoustic film (polyvinyl chloride) attached to the bottom [see Fig. 1(a) and (b)]. 3-D printing was used to fabricate the clip-on housing device frame using polylactic acid (PLA). The net weight of the device frame was 26 grams, and the total weight, including both reflectors, was 57 grams. An adapter was 3-D printed to allow a GE 9LD transducer (GE Healthcare, Wauwatosa, WI) to be easily attached to or removed from the FASTER device (Fig. 1(a)). The detailed design parameters are reported in Table I.

**Table I.** Specification of the FASTER Device Designed for the GE 9LD Linear Array Transducer

| Parameter | Value |
| --- | --- |
| Clip-on housing device weight (including reflectors) [g] | 57 |
| Clip-on housing device size ($l \times w \times h$) [mm<sup>3</sup>] | $74.8 \times 59.8 \times 30.9$ |
| Redirecting reflector size ($l \times w \times h$) [mm<sup>3</sup>] | $66.5 \times 17.0 \times 0.2$ |
| Fast-tilting reflector size ($l \times w \times h$) [mm<sup>3</sup>] | $48.4 \times 7.0 \times 0.2$ |
| Electromagnet size [mm<sup>2</sup>] | 10.0 diameter × 25.0 |
| Acoustic window size [mm<sup>2</sup>] | 50.9 lateral × 12.0 elevational |
| Acoustic film thickness [mm] | 0.01 |
| Power [W] | 0.48 |

Similar to the imaging procedures introduced in our previous work [27], the 1-D ultrasound transducer and the fast-tilting reflector were synchronized by triggering the sinusoidal driving signal of the fast-tilting reflector with the ultrasound imaging system. Corresponding tilting angles (e.g., *θ* in Fig. 1(c)) at each elevational location were calibrated. Because the incident angle is equal to the reflection angle of an acoustic beam, the final elevational scanning angle is 2*θ* after passing through two reflectors (Fig. 1(c)). The raw ultrasound data sampled in polar coordinates were scan converted to Cartesian coordinates using 3-D natural neighbor interpolation [28].

### Acoustic Validation Study of Ultrasound Beam Reflections

To evaluate the accuracy and efficacy of beam reflection by the redirecting reflector and the fast-tilting reflector, the acoustic field was carefully scanned and characterized using an acoustic intensity measurement system (AIMS III, Onda Corp, Sunnyvale, CA) with a capsule hydrophone (HGL-0200, Onda Corp, Sunnyvale, CA). 3-D acoustic pressure fields were measured in two different setups, as shown in Fig. 2: a) direct measurements from the ultrasound transducer without the FASTER device; and b) measurements with the FASTER device attached to the probe. For all the experiments, a Verasonics Vantage 256 system (Verasonics Inc., Kirkland, WA) was used. The hydrophone was mounted on the linear and rotary positioners of the ONDA AIMS system to scan the 3-D acoustic field, with 0.2-, 1-, and 2-mm step size in the elevational, axial, and lateral dimensions, respectively. The Vantage system was synchronized with the capsule hydrophone and the scanning stage.

**Fig. 2.**
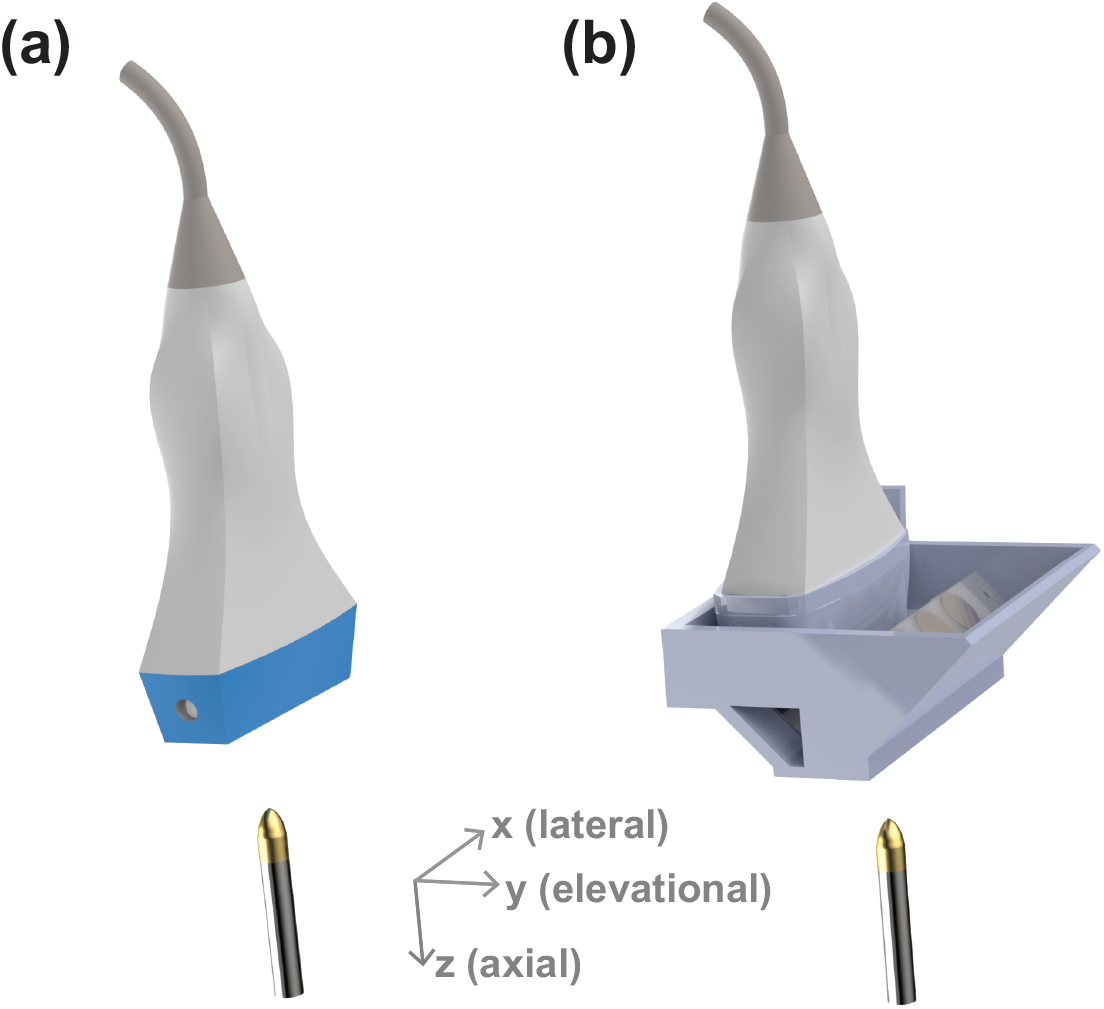
Experimental setup of the acoustic validation study for the FASTER 3-D imaging device. A hydrophone was used to measure the 3-D acoustic field without (a) and with (b) the FASTER device. For the measurements with the FASTER device attached, the fast-tilting reflector was kept stationary during the experiment.

The hydrophone-measured signals were processed using low-pass filtering and windowing. The acoustic beam characteristics (i.e., beamwidth and spectrum) were compared to examine if any beam distortion occurred inside the FASTER device. The beamwidth was quantified using the Full Width at Half Maximum (FWHM).

### Acoustic Calibration Study for Ultrasound Beam Scanning

To accurately reconstruct the volumetric images based on individual 2-D slices acquired at different scanning positions, the actual scanning angle of each 2-D imaging slice needs to be determined. Similar to the calibration process presented in the previous study [27], a second 1-D linear transducer (L7-4, ATL Philips, WA) was used to capture the dynamic 3-D scanning field of the FASTER 3-D imaging device, as shown in Fig. 3. The two 1-D transducers were synchronized with the FASTER device. The L7-4 transducer was mounted on the same linear and rotary positioners to scan the 3-D field, thus the 4-D (3-D space and time) scanning field can be accurately obtained. The sweeping motion of the transmitted beam was fitted using a sinusoidal function to match the sinusoidal driving signal of the fast-tilting reflector. By fitting the dynamic sweeping beams at different depths, the scanning characteristics (i.e., amplitude, initial phase, and offset) of the fast-tilting reflector were calibrated, enabling accurate volumetric reconstruction using the FASTER 3-D imaging device.

**Fig. 3.**
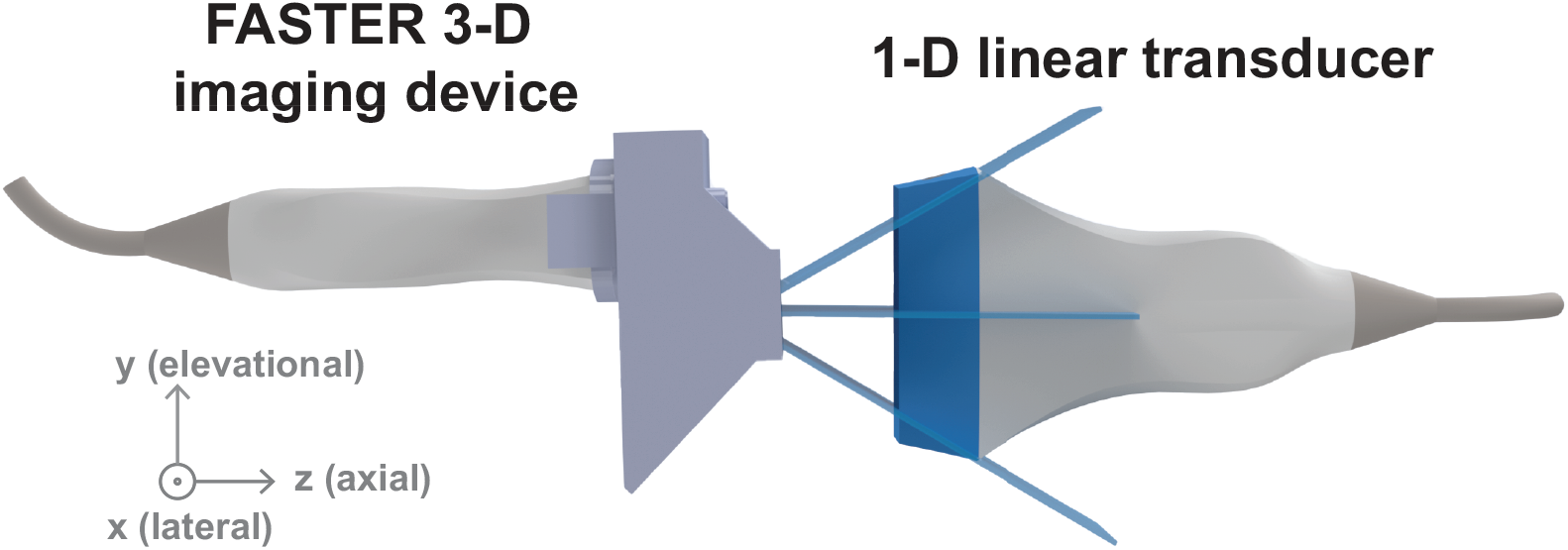
Experimental setup of the acoustic calibration study for the FASTER 3-D imaging device. A second 1-D linear transducer (L7-4) was used to measure the dynamic 4-D scanning field of the FASTER device.

### In Vitro Volumetric Imaging Studies

#### 3-D B-mode Imaging

To quantitatively evaluate the imaging performance of the new FASTER device, a wire phantom (tungsten 99.95%, 100 65 diameter, 4 wires, and 3 mm spacing) and a home-made tissue-mimicking phantom with an anechoic cylindrical cyst were imaged by the FASTER device attached to the GE 9LD transducer. Spatial resolution and contrast-to-noise ratio (CNR) were used as quantitative metrics for the evaluation. The frequency of the fast-tilting reflector was 150 Hz, resulting in a volume rate of 300 Hz as each scanning position was passed twice during one scanning cycle. For benchmarking, a mechanical translation-based 3-D image was captured to mimic that from a wobbler probe. The mechanical translation was precisely controlled by using a positing system (Daedal, Inc., Harrison City, PA, USA). A 0.5-mm step size was used along the elevational direction for the mechanical translation. The scanning volume rate was limited to approximately 0.1 Hz due to the scanning speed limitation of the Daedal system. The spatial resolution was characterized using the FWHM measured from the cross-sectional profiles of the wire targets. Fig. 4 illustrates the imaging setups, and the imaging parameters are summarized in Table II.

**Fig. 4.**
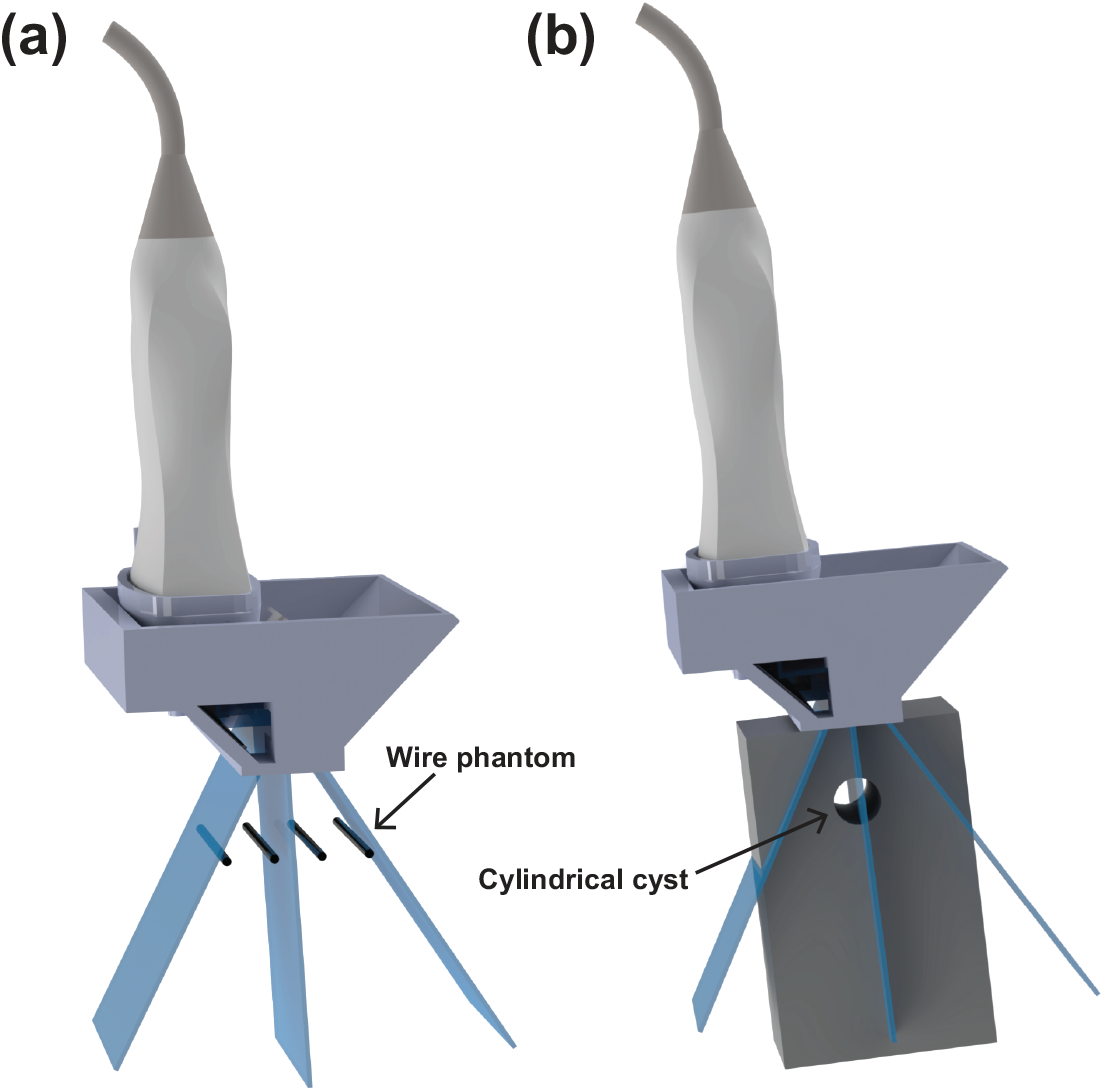
Experimental setup of the wire phantom imaging study (a) and the tissue-mimicking phantom imaging study (b) using the FASTER 3-D imaging device.

**Table II.** FASTER 3-D Imaging Parameters

| Parameter | Value |
| --- | --- |
| Central frequency [MHz] | 5.2 |
| Bandwidth [%] | 75 |
| Number of elements | 192 |
| PRF [Hz] | 9000 |
| Volume Rate [Hz] | 300 |
| Fast-tilting reflector tilting frequency [Hz] | 150 |
| Scanning range [°] | $\pm 24.8$ |
| Distance from the fast-tilting reflector to the transducer [mm] | 23.8 |
| Imaging depth [mm] | 60 |
| Lateral (x) range [mm] | 44.2 |
| Elevational (y) range [mm] | 33.5 |

#### 3-D Power Doppler Imaging

An *in vitro* 3-D PD study was performed using a cross-shaped flow phantom (1.48 mm diameter) with flowing microbubbles (Definity, concentration of 1.2 × 10^6^ MBs/mL, Lantheus Medical Imaging, Inc.). The detailed procedures of flow phantom making can be found in our previous work [29]. A constant flow velocity of 20 mm/s was used in this experiment. The imaging configurations were similar to the previous B-mode phantom study except that the frequency of the fast-tilting reflector was changed to 125 Hz with a PRF of 8000 Hz, which is equal to a 250 Hz volume rate. A total of 500 volumes were acquired with a 2-second acquisition time. Clutter filtering based on the singular value decomposition (SVD) [30] was performed on the reconstructed in-phase and quadrature (IQ) volumes, which were integrated to construct the 3-D PD image.

#### 3-D Super-Resolution Ultrasound Localization Microscopy

To further demonstrate the high volume rate imaging capability of the FASTER device, an *in vitro* 3-D ULM study was performed using the same cross-shaped flow phantom and imaging configurations as the 3-D PD study. A total of 5500 volumes were acquired and beamformed, followed by SVD clutter filtering and 3-D microbubble localization using the radial symmetry method [31, 32]. Finally, the trajectories of paired microbubbles across volumes were retrieved using a 3-D particle tracking algorithms [33] and accumulated into the final 3-D ULM intensity map. The 3-D flow velocity map was also calculated based on the microbubble trajectories. A voxel size of 46 65 along three dimensions was used for the ULM intensity and velocity maps.

#### In Vivo Volumetric Imaging Study

An *in vivo* study was performed on the basilic vein of a healthy volunteer. The same clinical GE 9LD ultrasound transducer with the FASTER device was used for imaging, and the imaging configurations were the same as the *in vitro* 3-D B-mode study (Table II). To accommodate the speed of sound difference between acoustic medium and human tissue (e.g., 1480 m/s vs 1540 m/s), the volumetric image was reconstructed using a customized delay-and-sum (DAS) beamforming, in which the delay was calculated using the Fast Marching method (FMM) by modeling a two-layer speed of sound map [34].

## Results

### Acoustic Validation Study of Ultrasound Beam Reflections

Fig. 5 shows the acoustic beam characterizations between the reference acoustic field (without the FASTER device) and the acoustic field with the FASTER device. The acoustic field from the FASTER device was close to the reference acoustic field [see Fig. 5(a) and (b)] without significant distortion from beam reflections. The elevational beam profile, as well as the beamwidth of the FASTER device, remained similar to the reference. For example, the FWHM of the reference acoustic field was 2.89 mm at 50-mm axial depth, and that of the FASTER device was 3.02 mm, as shown in Fig. 5(c). Furthermore, the spectrum of the acoustic field was consistent between the reference and FASTER device, as shown in Fig. 5(d). The beam characterizations show that the reflectors did not significantly distort the acoustic signals in the context of beamwidth and frequency response. This result can be attributed to the high reflection coefficient of the silicon wafer and the precise alignment between the ultrasound probe and the reflectors [see Fig. 1(c)]. Such precise alignment was made possible by the careful assembly of the clip-on housing device.

**Fig. 5.**
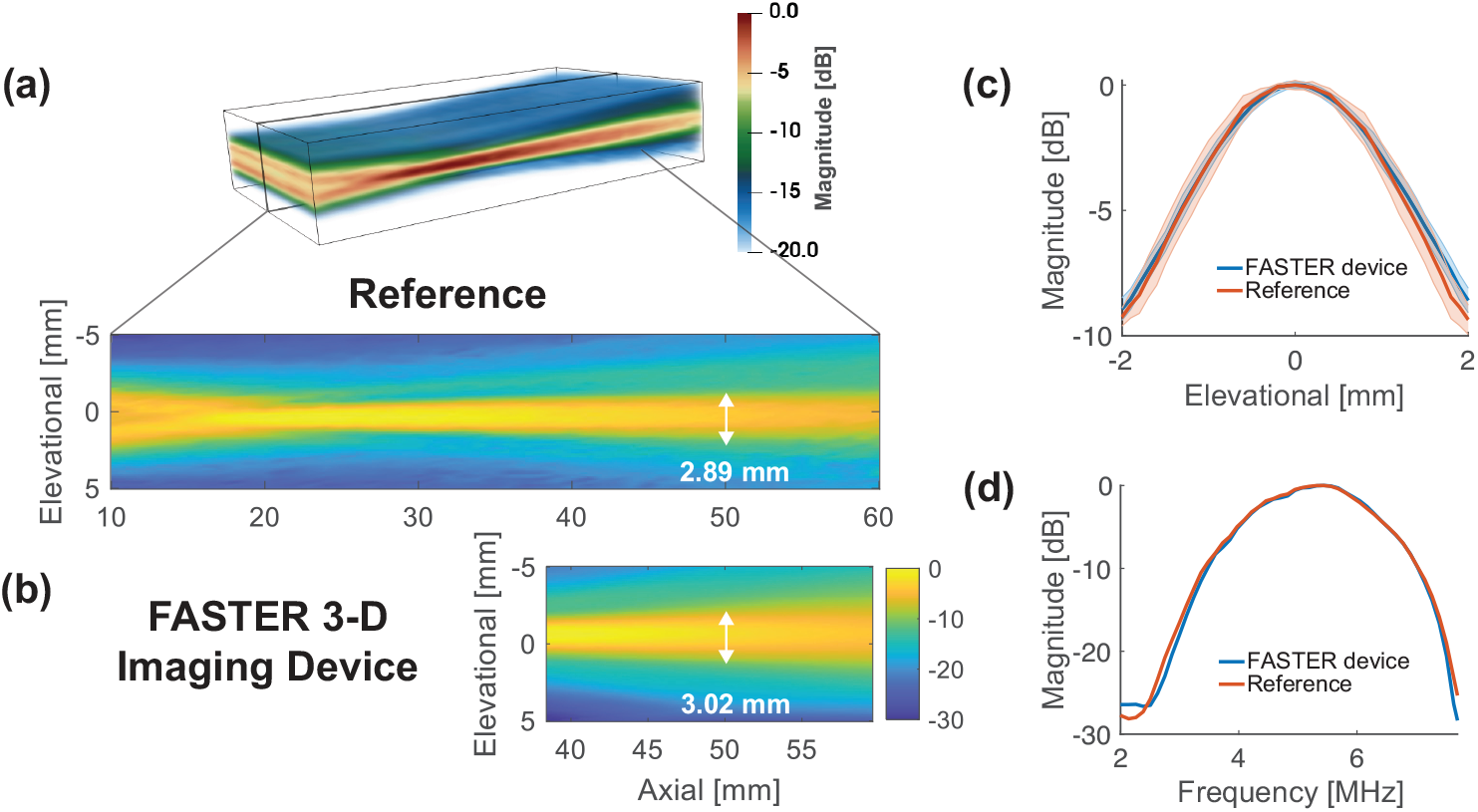
Acoustic validation study results for the FASTER 3-D imaging device. (a) Representative beam profile plots in the elevational-axial plane for the direct acoustic field measurement from the transducer (without FASTER device) and (b) the acoustic field of the FASTER device after the reflections by the redirecting reflector and fast-tilting reflector. The elevational beamwidths at 50 mm depth are labeled. (c) Plots of the beam profiles along the elevational dimension at 50-mm depth for reference and the FASTER device. (d) Comparisons of the spectrum of the acoustic fields with and without (reference) the FASTER device.

### Acoustic Calibration Study for Ultrasound Beam Scanning

Fig. 6 shows acoustic calibration results, including two transmitted beams measured at 36- and 45-mm axial depth over time. The sweeping motion of the transmitted beam was well matched with the sinusoidal driving signal of the fast-tilting reflector (red fitted curve). As illustrated in Fig. 6, through using the fitted sweeping motion measured at different axial locations, the scanning angle of the fast-tilting reflector was estimated as ±24.8 degrees with an initial position at -0.2 degree.

**Fig. 6.**
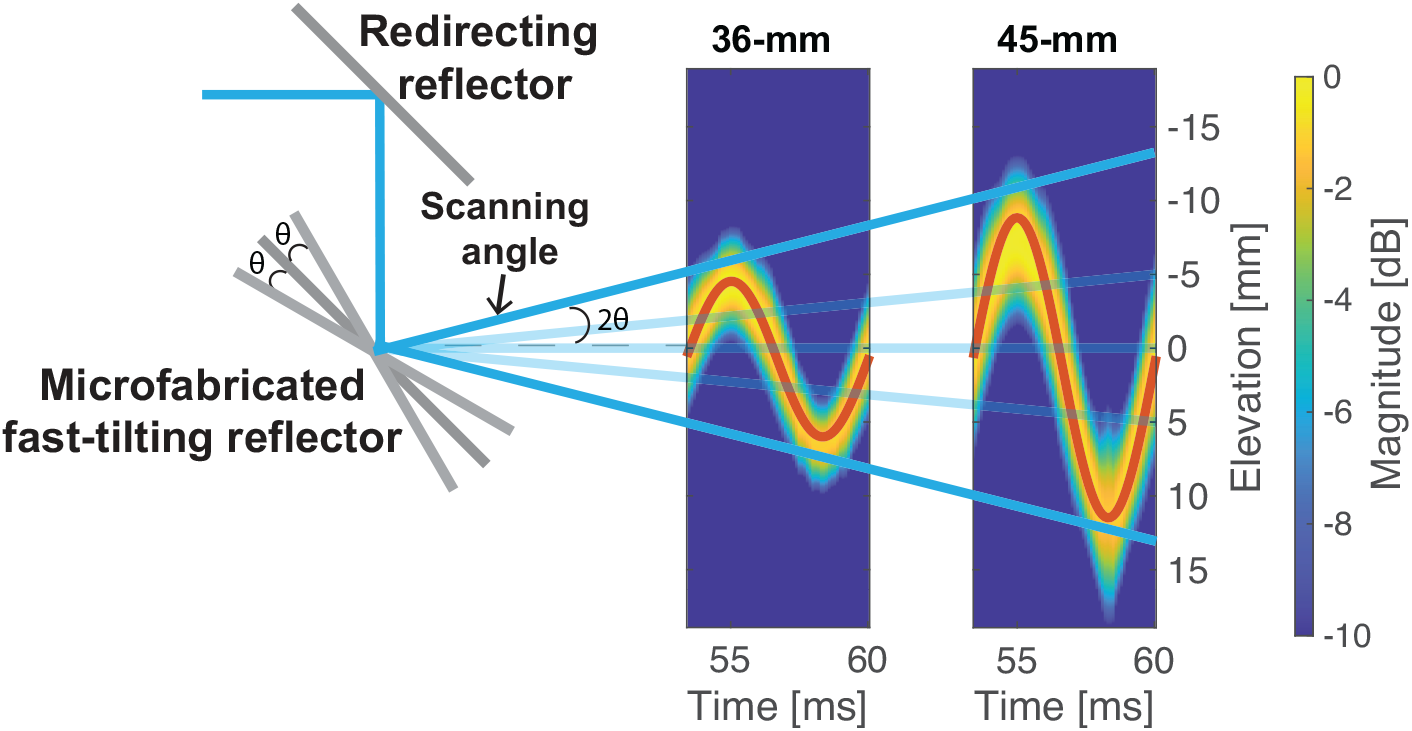
Acoustic calibration study results for the FASTER 3-D imaging device. Acoustically measured elevational beam intensities were plotted over time at 36- and 45-mm depths. Low-pass filtering and windowing were applied to the raw radio-frequency (RF) data acquired by the L7-4 transducer, and sinusoidal fitting was used on the beam intensity over time to characterize the dynamic scanning of the FASTER device.

### In Vitro Volumetric Imaging Studies

#### 3-D B-mode Imaging

Fig. 7 shows the wire phantom imaging results using the FASTER 3-D imaging device and the simulated wobbler 3-D scanning based on mechanical translation. The four wires imaged using the FASTER device were accurately reconstructed and clearly isolated from each other. The image from the FASTER device shows good agreement with that from the mechanical translation. The elevational FWHM of the wire imaged by the FASTER device was 1.73 ±0.06 mm, and the one imaged by the linear scanning method was 1.93 ±0.19 mm. The wire phantom imaging results demonstrated that the proposed FASTER device using both redirecting and fast-tilting reflectors provides comparable elevational resolution to the reference wobbler scanning method but with much higher volume rates (e.g., 300 Hz for FASTER vs. ∼0.1 Hz for mechanical translation).

**Fig. 7.**
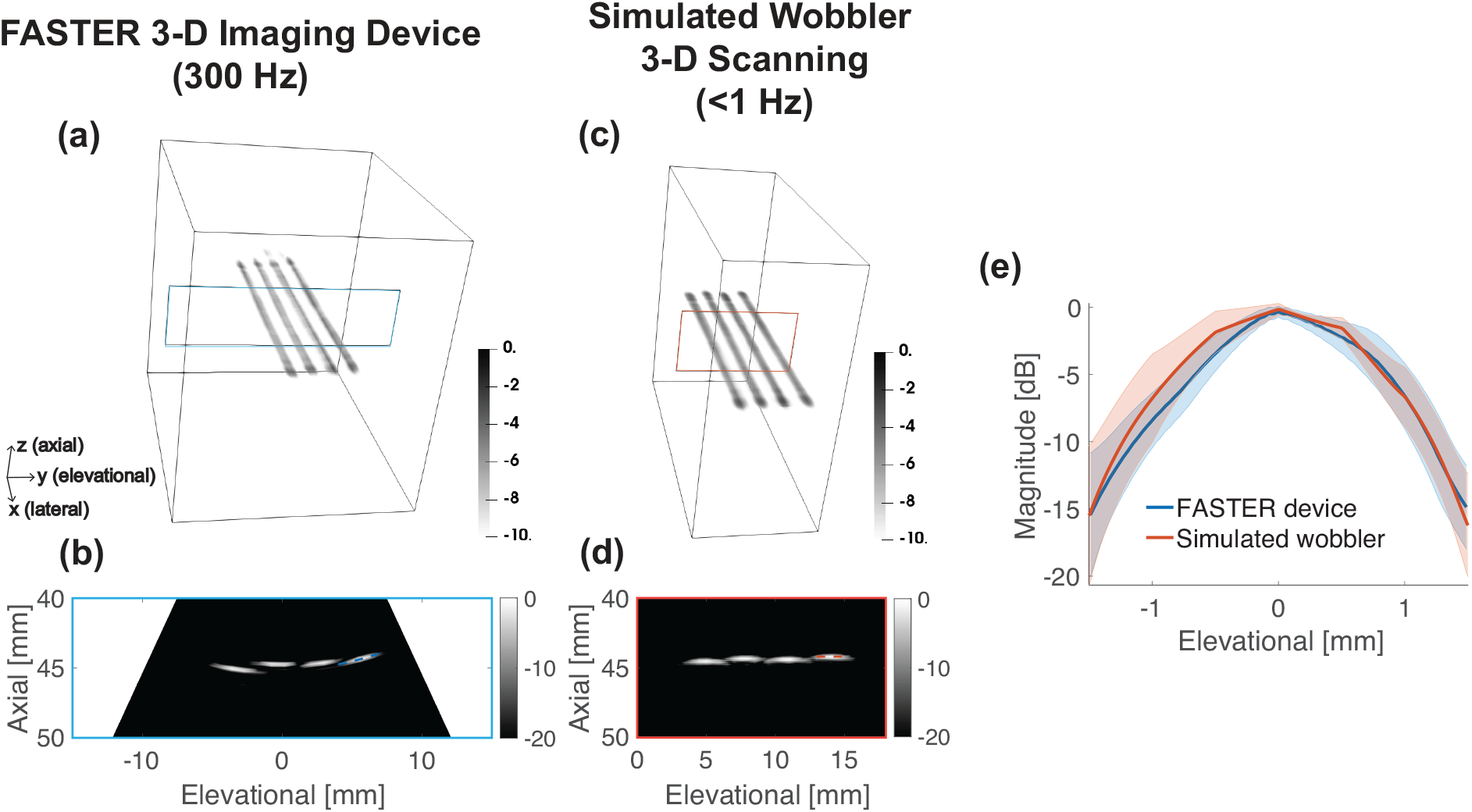
Wire phantom imaging results using the FASTER 3-D imaging device and the simulated wobbler. (a) Reconstructed volumetric image of the wire phantom using the FASTER 3-D imaging device at 300 Hz volume rate. (b) Elevational-axial image of the wire phantom using the FASTER 3-D imaging device. (c) Reconstructed volumetric image of the wire phantom using the simulated wobbler. (d) Elevational-axial image of the wire phantom using the simulated wobbler scanning. (e) Elevational profiles of the wire imaged by the FASTER 3-D imaging device and the simulated wobbler probe, corresponding ROIs are labeled in (b) and (d), and standard deviation (shaded area) was calculated across different lateral locations.

Fig. 8 shows the tissue-mimicking phantom results using the FASTER 3-D imaging device and the simulated wobbler 3-D scanning. From both the 3-D volumetric rendering and 2-D images in

**Fig. 8.**
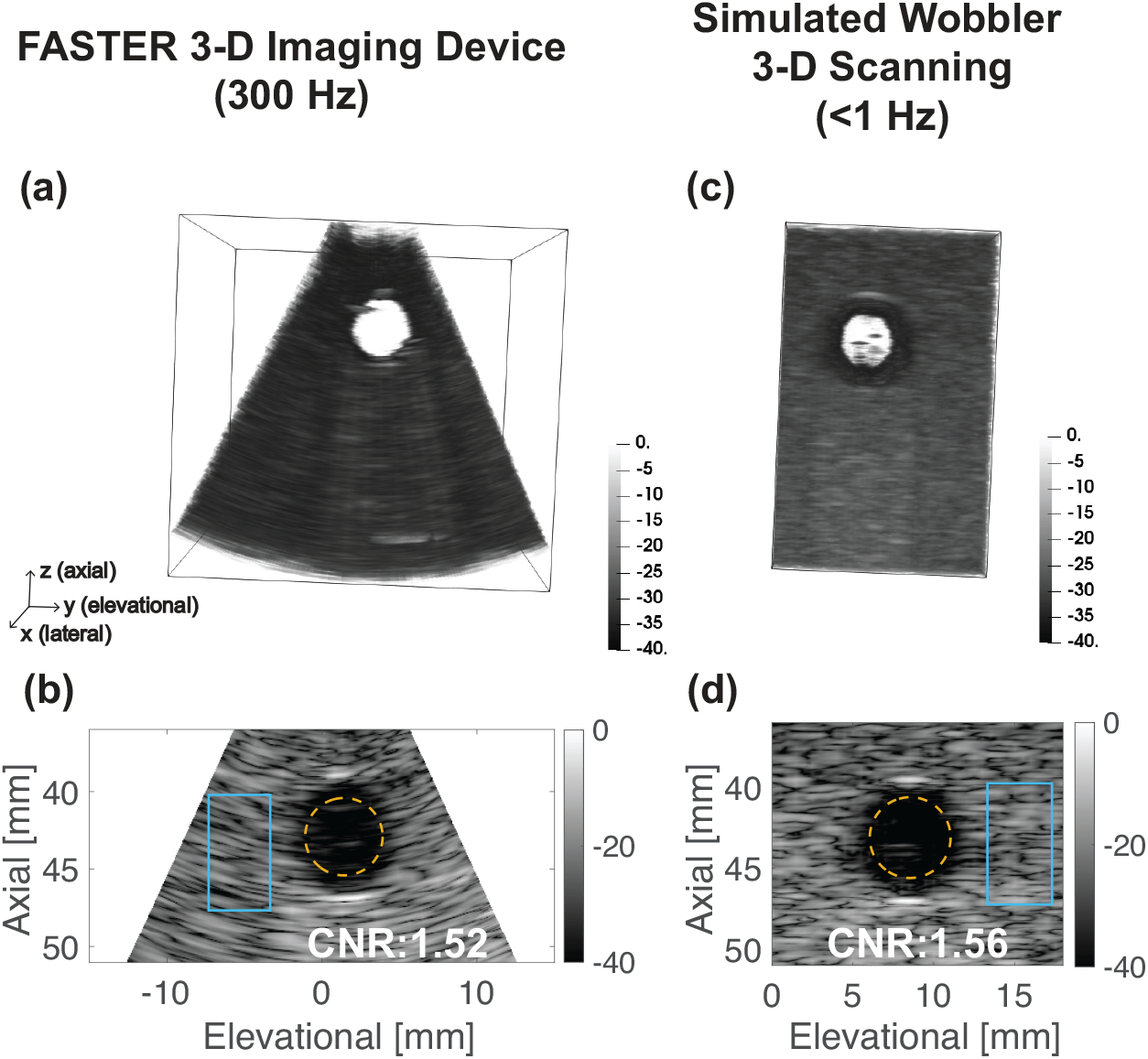
Tissue-mimicking phantom imaging results using the FASTER 3-D imaging device and simulated wobbler 3-D scanning. (a) Reconstructed volumetric image of the tissue-mimicking phantom using FASTER 3-D imaging device at 300 Hz volume rate. (b) Elevational-axial image of the tissue-mimicking phantom using the FASTER 3-D imaging (c) Reconstructed volumetric image of the tissue-mimicking phantom using simulated wobbler 3-D scanning. (d) Elevational-axial image of the tissue-mimicking phantom using simulated wobbler 3-D scanning. The ROIs for CNR calculation are labeled in (b) and (d) with calculated CNR values.

Fig. 8, we can clearly see an anechoic cyst reconstructed with high contrast to the background using both methods. The CNR of the anechoic cyst imaged by the FASTER device was 1.52, and the reference CNR value by the scanning method was 1.56. The cyst-like inclusion phantom results showed that the FASTER device could provide similar imaging performance in the context of contrast to the benchmark method.

#### 3-D Power Doppler Imaging

Fig. 9(a) and (b) show the reconstructed PD image of the cross-shaped flow phantom using the FASTER 3-D imaging device, where the two flow channels that are orthogonal to each other are clearly perceived. The measured lateral FWHM of the cross-section of the flow channel located at 45-mm depth was 1.47 mm, as shown in Fig. 9(g). The 3-D PD imaging study illustrated the potential capability of contrast-free blood flow imaging with the FASTER device.

**Fig. 9.**
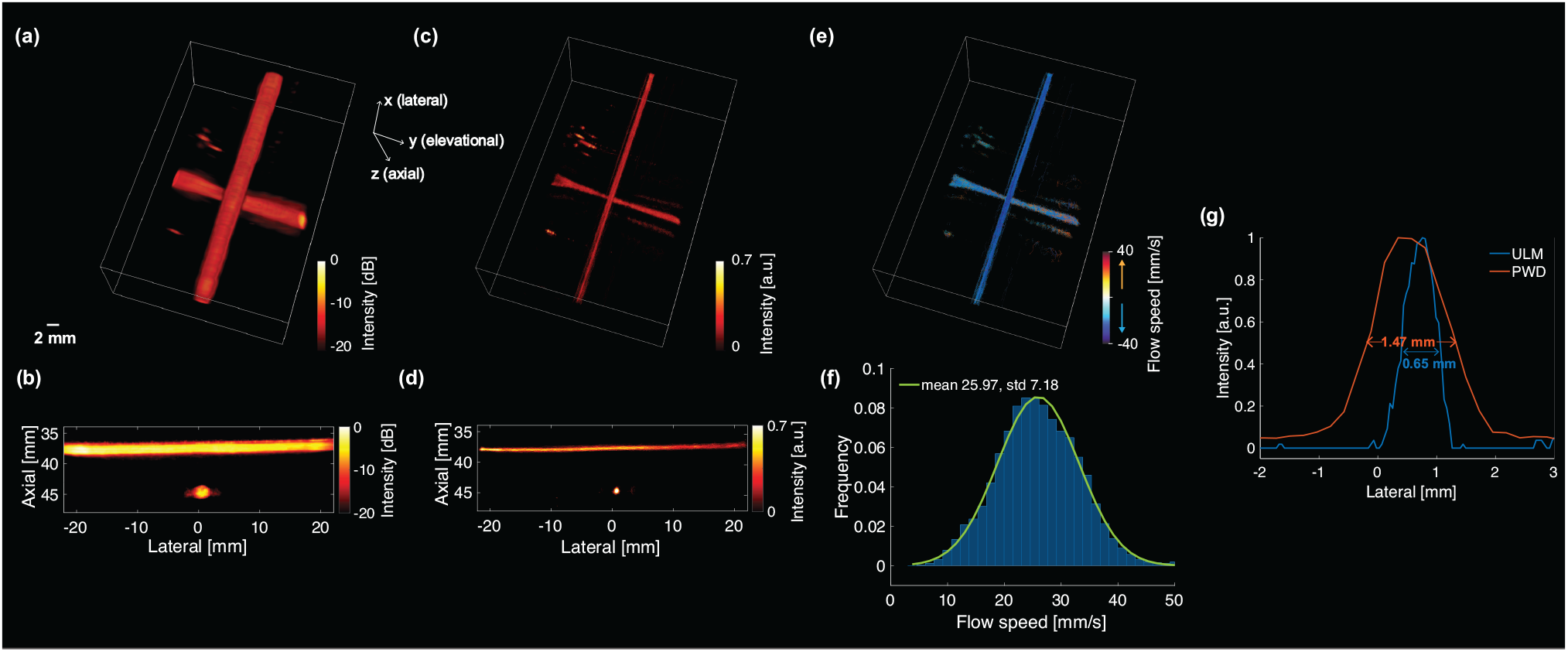
3-D PD imaging and 3-D super-resolution ULM imaging of the cross-shaped flow phantom using the FASTER 3-D imaging device. (a) Reconstructed volumetric PD image of the flow phantom. (b) Lateral-axial PD image. (c) Reconstructed volumetric ULM intensity map of the flow phantom. The map was compressed with a cubic root for enhanced visibility. (d) Lateral-axial ULM intensity map (1-mm thickness and summation projection). (e) Reconstructed volumetric ULM flow velocity map of the flow phantom. (f) Flow velocity histogram of the flow phantom with a gaussian fitted curve (mean is 25.97 mm/s, standard deviation is 7.18 mm/s). (g) PD and ULM intensity profiles along the lateral dimension of the flow channel located at 45-mm depth.

#### 3-D Super-Resolution Ultrasound Localization Microscopy

Fig. 9(c)-(e) show the reconstructed ULM intensity map and flow velocity map of the cross-shaped flow phantom using the FASTER 3-D imaging device. The two perpendicular flow channels were clearly reconstructed with flow velocity estimation at a much higher spatial resolution. The mean velocity across all voxels was 25.97 ±7.18 mm/s, which was higher than the nominal flow speed setup (20 mm/s), as shown in Fig. 9(f). One possible reason is that the gelatin-made flow channels may have been compressed slightly during acoustic coupling, resulting in a reduced luminal diameter and a higher flow speed inside the channels. The measured lateral FWHM of the same flow channel located at 45-mm depth was 0.65 mm (Fig. 9(g)), which was more than 2-fold improvement compared to the 3-D PD imaging results. However, it should be noted that FWHM may not be an appropriate metric for ULM to measure the flow channel diameter due to the stochastic nature of microbubble trajectory reconstruction. The 3-D ULM imaging study demonstrated the promising capability of applying advanced imaging techniques that need high-speed 3-D motion tracking with the FASTER device.

### In Vivo Volumetric Imaging Study

Fig. 10 shows the *in vivo* imaging results using the FASTER 3-D imaging device on a healthy volunteer. The basilic vein was imaged, which can be clearly observed from the elevational-axial plane (cross-sectional view) and in the lateral-axial plane (longitudinal view) (arrows in Fig. 10). By providing volumetric data, any cross-sectional slicing of the vessel can be achieved, allowing for the quantification of physiological biomarkers such as vessel diameter and volume with anatomical context. A real-time bi-planar B-mode imaging mode (Fig. 10(c)) provides a convenient and computationally efficient way of displaying 3-D volumetric data [35]. These results demonstrate the capability of *in vivo* human imaging using the FASTER device with a high volumetric imaging rate (e.g., 300 Hz).

**Fig. 10.**
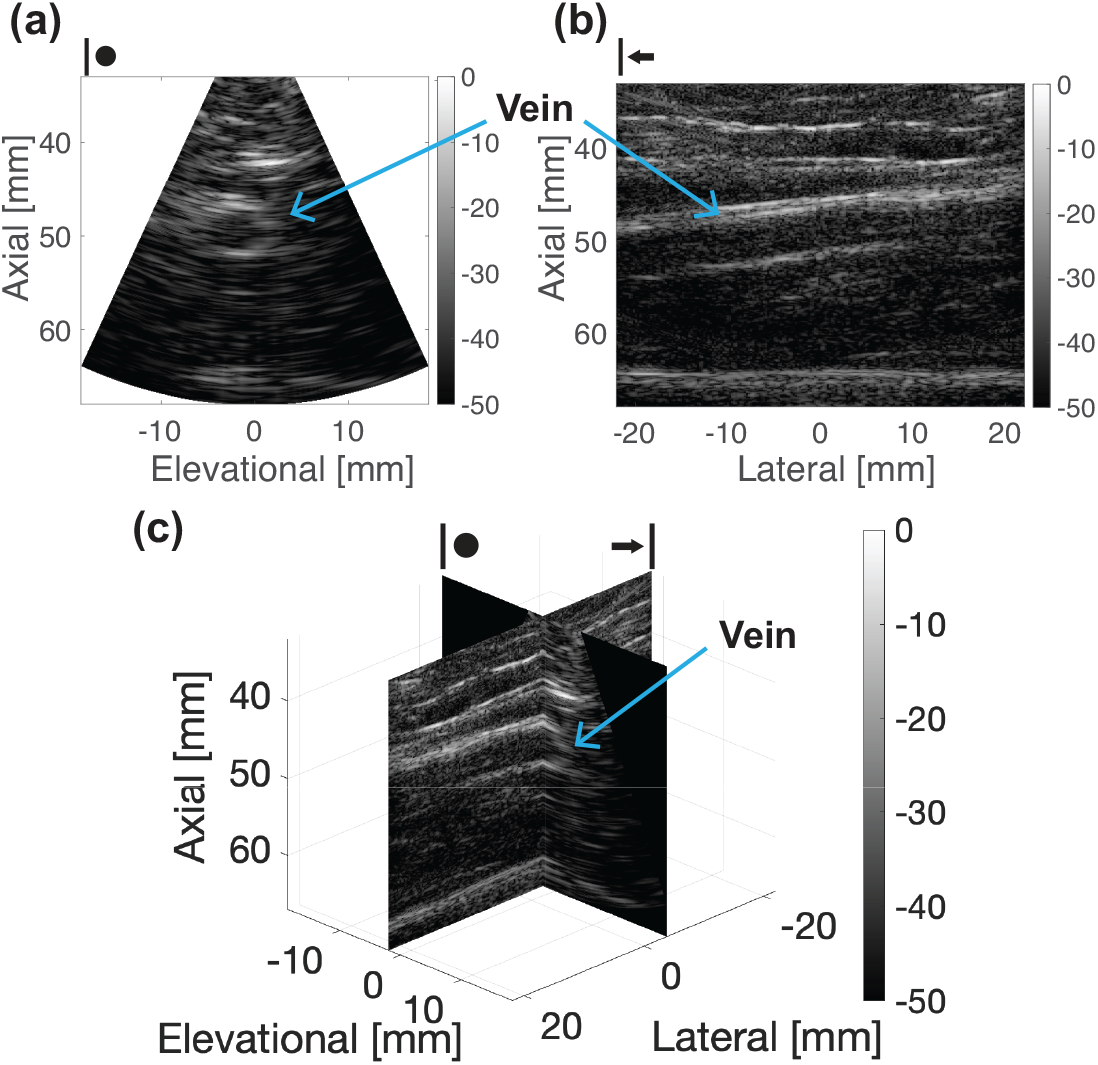
*In vivo* 3-D imaging of the basilic vein from a healthy volunteer using the FASTER 3-D imaging device. (a) Out-of-plane (elevational-axial) image of the vein. (b) In-plane (lateral-axial) image of the vein. (c) A bi-plane view of the vein.

## Discussions

Currently, 3-D ultrasound imaging is challenged by various technical limitations, such as low volume rate, high cost of equipment, and high computational complexities involved with beamforming and post-processing. To address these issues, here we proposed a novel 3-D imaging technique with high volumetric imaging rate – FASTER – which uses a fast-tilting reflector and a redirecting reflector to rapidly sweep plane waves in the elevational direction for high-speed 3-D scanning. Our results demonstrate that FASTER could achieve a high imaging volume rate (e.g., hundreds of Hertz) and comparable imaging quality as compared to conventional mechanical-translation-based 3-D scanning (e.g., wobbler). Additionally, FASTER is based on conventional 1-D array transducers and can be conveniently adapted to existing ultrasound transducers. FASTER also enjoys a lightweight and compact form factor that is user-friendly. These advantages of FASTER pave the way for future clinical translations of the technology.

The previous FASTER solution changed the incident beam direction 90 degrees (i.e., from axial to elevational), which makes it challenging to perform hand-held scanning. In this article, we improved the FASTER technique by incorporating a redirecting reflector to recover the original incident beam direction, which allows the transducer to be held in a regular position (e.g., upright). The validation study demonstrated that there were no significant beam distortions introduced by the new redirecting reflector.

Another limitation of the previous FASTER technique was that the imaging had to be performed inside a water tank because of the need of water as an acoustic wave conducting medium, which is not a practical solution for *in vivo* imaging. To address this issue, a clip-on housing device was designed to enclose the acoustic medium together with the redirecting reflector and the fast-tilting reflector. The clip-on device can be conveniently attached to or removed from conventional 1-D ultrasound probes. To avoid leakage of the acoustic medium, an acoustic film was sealed on the bottom of the device. The clip-on housing device frame can be easily fabricated using 3-D printing with biocompatible materials at a low cost, and the entire clip-on device is lightweight (57 grams).

The *in vitro* phantom imaging studies showed that the newly developed FASTER 3-D imaging device provided comparable spatial resolution and contrast while maintaining a large FOV (e.g., 50-degree scanning angle) and a high volume rate (e.g., 300 Hz). The *in vivo* study showed the possibility of using the FASTER device for *in vivo* human imaging. Multiple imaging modalities, including B-mode, PD, and ULM, were demonstrated. The flow phantom study showcased the potential of applying advanced imaging techniques (3-D ULM, and possibly 3-D SWE) using the new FASTER device. These results suggest that FASTER not only provides a viable solution for accessible 3-D ultrasound imaging, but it also has the potential to maintain and extend the functionality of 2-D ultrasound imaging to 3-D.

The current implementation of FASTER 3-D imaging requires synchronization between the tilting reflectors and the ultrasound system for accurate image reconstruction. For easier translation of the FASTER 3-D technology, an asynchronous acquisition mode is also feasible by using the correlation between consecutively acquired frames to estimate the scanning beam position. For example, the correlation coefficient between adjacent 2-D frames is the highest when the scanning angle is the largest (scanning speed is the lowest), and *vice versa*. The asynchronous acquisition capability should greatly reduce the burden of potential system modifications to allow FASTER to be implemented on existing commercial ultrasound systems.

There are many other advanced ultrasound imaging techniques that can be benefited from high volumetric imaging rates enabled by the FASTER device. For example, 3-D SWE is essential for accurate and comprehensive tissue stiffness measurement. However, 3-D SWE (acoustic radiation force-based) is difficult to achieve because of the need for a high 3-D imaging volume rate and transmitting the high-power push beams. Wobblers do not provide an adequate 3-D volume rate for shear wave tracking, while 2-D matrix arrays do not typically support the transmission of long-duration, high mechanical index (MI) push pulses for shear wave generation. In comparison, FASTER provides a potentially viable solution for 3-D SWE because it supports high volume rate 3-D scanning and it uses conventional 1-D ultrasound transducers that are compatible with ARF-based SWE. Although the volume rate of the current FASTER device (e.g., 300 Hz) is not adequate for 3-D SWE, ongoing studies are being conducted to increase the tilting frequency of the reflector for 3-D shear wave tracking (e.g., to 2 kHz).

The acoustic film in the current device design was made with Polyvinyl chloride (PVC) with a thickness of 0.01 mm. The acoustic impedance mismatch between the PVC acoustic film and the acoustic conducting medium (e.g., water) may result in reverberation artifacts for FASTER. Future work will be conducted to fabricate acoustic films using biocompatible materials with reduced reverberation and membrane thickness.

In addition to the impedance mismatch between the acoustic film and the acoustic medium, there was a slight speed of sound mismatch between the acoustic medium and soft tissue for *in vivo* imaging. To address this issue, a two-layer speed of sound map was used in the DAS beamforming process based on the Fast Marching method [34] for the *in vivo* imaging study. An alternative solution is to switch to an acoustic wave conduction medium with a similar speed of sound of soft tissue.

## Conclusions

In this work, we introduced an improved FASTER 3-D imaging method by incorporating a redirecting reflector to facilitate the convenient handling of the ultrasound probe. We also developed a compact and lightweight clip-on housing device that can be easily adapted to conventional 1-D transducers. The phantom imaging studies demonstrated that the FASTER 3-D imaging device provided comparable imaging quality to the conventional, mechanical-translation-based 3-D imaging (i.e., wobbler) while providing a much higher 3-D imaging volume rate. The utility of the new FASTER device was also tested in an *in vivo* case study for 3-D B-mode imaging. The proposed methods in this study could clear key hurdles for the application of FASTER in regular ultrasound imaging settings for high speed and high quality 3-D ultrasound imaging.

## Acknowledgment

This work was supported by the National Institute of Biomedical Imaging and Bioengineering of the National Institutes of Health under Award Number R01EB031040. The content is solely the responsibility of the authors and does not necessarily represent the official views of the National Institutes of Health.

## References

[1] G. Montaldo, M. Tanter, J. Bercoff, N. Benech, and M. Fink, “Coherent plane-wave compounding for very high frame rate ultrasonography and transient elastography,” IEEE transactions on ultrasonics, ferroelectrics, and frequency control, vol. 56, no. 3, pp. 489–506, 2009.

[2] M. Tanter and M. Fink, “Ultrafast imaging in biomedical ultrasound,” IEEE transactions on ultrasonics, ferroelectrics, and frequency control, vol. 61, no. 1, pp. 102–119, 2014.

[3] J. Bercoff, M. Tanter, and M. Fink, “Supersonic shear imaging: a new technique for soft tissue elasticity mapping,” IEEE transactions on ultrasonics, ferroelectrics, and frequency control, vol. 51, no. 4, pp. 396–409, 2004.

[4] P. Song et al., “Two-dimensional shear-wave elastography on conventional ultrasound scanners with time-aligned sequential tracking (TAST) and comb-push ultrasound shear elastography (CUSE),” IEEE transactions on ultrasonics, ferroelectrics, and frequency control, vol. 62, no. 2, pp. 290–302, 2015.

[5] E. Macé, G. Montaldo, I. Cohen, M. Baulac, M. Fink, and M. Tanter, “Functional ultrasound imaging of the brain,” Nature methods, vol. 8, no. 8, pp. 662–664, 2011.

[6] C. Errico et al., “Ultrafast ultrasound localization microscopy for deep super-resolution vascular imaging,” Nature, vol. 527, no. 7579, pp. 499–502, 2015.

[7] N. Renaudin, C. Demené, A. Dizeux, N. Ialy-Radio, S. Pezet, and M. Tanter, “Functional ultrasound localization microscopy reveals brain-wide neurovascular activity on a microscopic scale,” Nature methods, vol. 19, no. 8, pp. 1004–1012, 2022.

[8] R. W. Gill, “Measurement of blood flow by ultrasound: accuracy and sources of error,” Ultrasound in medicine & biology, vol. 11, no. 4, pp. 625–641, 1985.

[9] J.-l. Gennisson et al., “4-D ultrafast shear-wave imaging,” IEEE transactions on ultrasonics, ferroelectrics, and frequency control, vol. 62, no. 6, pp. 1059–1065, 2015.

[10] G. Unsgaard, S. Ommedal, T. Muller, A. Gronningsaeter, and T. A. Nagelhus Hernes, “Neuronavigation by intraoperative three-dimensional ultrasound: initial experience during brain tumor resection,” Neurosurgery, vol. 50, no. 4, pp. 804–812, 2002.

[11] D. C. D. A. Bastos et al., “Challenges and opportunities of intraoperative 3D ultrasound with neuronavigation in relation to intraoperative MRI,” Frontiers in oncology, vol. 11, p. 656519, 2021.

[12] T. Lange et al., “3D ultrasound-CT registration of the liver using combined landmark-intensity information,” International journal of computer assisted radiology and surgery, vol. 4, no. 1, pp. 79–88, 2009.

[13] H. Neshat, D. W. Cool, K. Barker, L. Gardi, N. Kakani, and A. Fenster, “A 3D ultrasound scanning system for image guided liver interventions,” Medical physics, vol. 40, no. 11, p. 112903, 2013.

[14] M. V. Burri, D. Gupta, R. E. Kerber, and R. M. Weiss, “Review of novel clinical applications of advanced, real-time, 3-dimensional echocardiography,” Translational Research, vol. 159, no. 3, pp. 149–164, 2012.

[15] P. Acar et al., “Real-time three-dimensional foetal echocardiography using a new transabdominal xMATRIX array transducer,” Archives of Cardiovascular Diseases, vol. 107, no. 1, pp. 4–9, 2014.

[16] R. Chaoui, A. Abuhamad, J. Martins, and K. S. Heling, “Recent development in three and four dimension fetal echocardiography,” Fetal Diagnosis and Therapy, vol. 47, no. 5, pp. 345–353, 2020.

[17] A. Fenster, K. Surry, W. Smith, and D. B. Downey, “The use of three-dimensional ultrasound imaging in breast biopsy and prostate therapy,” Measurement, vol. 36, no. 3-4, pp. 245–256, 2004.

[18] M. Correia, J. Provost, M. Tanter, and M. Pernot, “4D ultrafast ultrasound flow imaging: in vivo quantification of arterial volumetric flow rate in a single heartbeat,” Physics in Medicine & Biology, vol. 61, no. 23, p. L48, 2016.

[19] R. W. Prager, U. Z. Ijaz, A. Gee, and G. M. Treece, “Three-dimensional ultrasound imaging,” Proceedings of the Institution of Mechanical Engineers, Part H: Journal of Engineering in Medicine, vol. 224, no. 2, pp. 193–223, 2010.

[20] J. T. Yen and S. W. Smith, “Real-time rectilinear 3-D ultrasound using receive mode multiplexing,” ieee transactions on ultrasonics, ferroelectrics, and frequency control, vol. 51, no. 2, pp. 216–226, 2004.

[21] S. Blaak et al., “Design of a micro-beamformer for a 2D piezoelectric ultrasound transducer,” in 2009 IEEE International Ultrasonics Symposium, 2009: IEEE, pp. 1338–1341.

[22] E. Roux, F. Varray, L. Petrusca, C. Cachard, P. Tortoli, and H. Liebgott, “Experimental 3-D ultrasound imaging with 2-D sparse arrays using focused and diverging waves,” Scientific reports, vol. 8, no. 1, pp. 1–12, 2018.

[23] C. E. Morton and G. R. Lockwood, “Theoretical assessment of a crossed electrode 2-D array for 3-D imaging,” in IEEE Symposium on Ultrasonics, 2003, 2003, vol. 1: IEEE, pp. 968–971.

[24] J. A. Jensen et al., “Three-dimensional super-resolution imaging using a row–column array,” IEEE transactions on ultrasonics, ferroelectrics, and frequency control, vol. 67, no. 3, pp. 538–546, 2019.

[25] J. Sauvage et al., “4D Functional imaging of the rat brain using a large aperture row-column array,” IEEE transactions on medical imaging, vol. 39, no. 6, pp. 1884–1893, 2019.

[26] Z. Dong et al., “Three-dimensional Shear Wave Elastography Using a 2D Row Column Addressing (RCA) Array,” BME Frontiers, vol. 2022, 2022.

[27] Z. Dong, S. Li, M. R. Lowerison, J. Pan, J. Zou, and P. Song, “Fast Acoustic Steering Via Tilting Electromechanical Reflectors (FASTER): A Novel Method for High Volume Rate 3-D Ultrasound Imaging,” IEEE transactions on ultrasonics, ferroelectrics, and frequency control, vol. 68, no. 3, pp. 675–687, 2020.

[28] I. Amidror, “Scattered data interpolation methods for electronic imaging systems: a survey,” Journal of electronic imaging, vol. 11, no. 2, pp. 157–176, 2002.

[29] X. Chen et al., “Localization free super-resolution microbubble velocimetry using a long short-term memory neural network,” bioRxiv, p. 2021.10. 01.462404, 2021.

[30] J. Baranger, B. Arnal, F. Perren, O. Baud, M. Tanter, and C. Demené, “Adaptive spatiotemporal SVD clutter filtering for ultrafast Doppler imaging using similarity of spatial singular vectors,” IEEE transactions on medical imaging, vol. 37, no. 7, pp. 1574–1586, 2018.

[31] B. Heiles et al., “Volumetric ultrasound localization microscopy of the whole rat brain microvasculature,” IEEE Open Journal of Ultrasonics, Ferroelectrics, and Frequency Control, vol. 2, pp. 261–282, 2022.

[32] R. Parthasarathy, “Rapid, accurate particle tracking by calculation of radial symmetry centers,” Nature methods, vol. 9, no. 7, pp. 724–726, 2012.

[33] K. Jaqaman et al., “Robust single-particle tracking in live-cell time-lapse sequences,” Nature methods, vol. 5, no. 8, pp. 695–702, 2008.

[34] J. A. Sethian, “Fast marching methods,” SIAM review, vol. 41, no. 2, pp. 199–235, 1999.

[35] D. Convissar, E. A. Bittner, and M. G. Chang, “Biplane Imaging Versus Standard Transverse Single-Plane Imaging for Ultrasound-Guided Peripheral Intravenous Access: A Prospective Controlled Crossover Trial,” Critical Care Explorations, vol. 3, no. 10, 2021.

